# Crossmodal plasticity effects of hearing loss and ageing in the human auditory cortex

**DOI:** 10.64898/2026.06.08.730968

**Authors:** Zheng Pan, Marcelina Połeć, Angelika Avgerinos, Krislynn Bi, Tim Green, Velia Cardin

## Abstract

Crossmodal plasticity refers to the phenomenon by which typical sensory areas respond to stimulation in other sensory modalities when their main input is absent or reduced. In this study, we examined the effect of age-related hearing loss (ARHL) on crossmodal plasticity in auditory brain regions. ARHL is prevalent in a large proportion of older adults, and it is associated with an increased risk of cognitive decline and dementia. Understanding how the brain adapts to ARHL is essential in aiding the development of adequate therapies and interventions for a phenomenon experienced by most adults aged 65 and over. Previous research in congenitally deaf adults has shown that crossmodal plasticity effects are stronger for conditions that require higher executive demands. We conducted an fMRI experiment in older adults to determine whether crossmodal recruitment of auditory cortical regions during higher executive demands also occurs following ARHL. In the MRI scanner, participants with and without ARHL completed a visual task-switching paradigm and a visual working memory task, each comprising high and low executive-demand conditions. Results from both groups of participants showed that the high-demand conditions reliably activated canonical frontoparietal regions involved in executive function and cognitive control. Crucially, we also observed significant recruitment of auditory regions during these visual tasks, particularly under the higher executive demands condition of the task-switching paradigm. Crossmodal activations in auditory areas occurred in both groups, with no significant effects of hearing level or age. These findings indicate that auditory regions are involved in high executive demand visual processing, and that crossmodal plasticity effects are not restricted to early sensitive periods. We propose that reduced auditory input and age-related changes in cortical processing jointly contribute to the reorganisation of the auditory cortex in later life. Understanding these mechanisms is critical for clarifying the neural consequences of ARHL and their impact on cognitive decline.

## Introduction

Hearing loss is highly prevalent in older age: globally, more than 60% of adults aged 65 years and above have a degree of hearing loss (World Health Organization, 2021). In the context of rapidly ageing populations and given its association with cognitive decline and dementia, ARHL represents a significant medical and socioeconomic challenge (Lin et al., 2013; Stucky et al., 2010; Uchida et al., 2019; Armstrong et al., 2020; Griffiths et al., 2020; Livingston et al., 2020; Shende & Mudar, 2023; Akeroyd & Munro, 2024; Loughrey, 2025).

ARHL commonly results from changes in peripheral auditory structures, which decrease the effectiveness in transduction and the neural decoding of sounds (Bowl & Dawson, 2019; Slade et al., 2020; Elliott et al., 2022). ARHL also has a significant impact on the structure and function of the brain, especially the central auditory system (Cardin, 2016; Slade et al., 2020; Herrmann & Butler, 2021). Neuroimaging studies have shown that ARHL results in functional changes in cortical auditory and speech processing networks (Peelle et al., 2011; Fitzhugh et al., 2019; Wolak et al., 2019; Huang et al., 2022). While there is significant evidence showing the effects of ARHL on auditory processing in cortical areas, it is less clear if ARHL also impacts crossmodal plasticity in these regions. Crossmodal plasticity refers to the phenomena by which typical sensory areas respond to stimulation in other sensory modalities when their main input is absent or significantly reduced (Bavelier & Neville, 2002; Sharma & Glick, 2016; Vinogradova & Cardin, 2024). The most striking cases of crossmodal plasticity in auditory cortices are found in congenitally deaf individuals, where regions of the brain typically involved in auditory processing respond to visual and tactile stimulation (Fine et al., 2005; Bottari et al., 2014; Karns et al., 2012; Almeida et al., 2015; Ding et al., 2015; Bola et al., 2017; Cardin et al., 2018; Zimmermann et al., 2021; Manini et al., 2022; Dhanik et al., 2024; Manini et al., 2025; Tal et al., 2026). Recent studies in congenitally and early deaf individuals have shown that crossmodal plasticity effects are not limited to sensory processing in other modalities. Research employing different cognitive tasks show that crossmodal responses are modulated by task demands, with auditory cortices in deaf individuals being more strongly activated by high-executive-demand cognitive tasks compared to perceptually matched control tasks with low executive demands (Ding et al., 2015; Cardin et al., 2018; Manini et al., 2022). These high-executive demand tasks typically elicit activity in frontoparietal regions. Studies in deaf individuals have shown that auditory brain regions, such as primary auditory cortex and posterior superior temporal cortex (pSTC), are also recruited during visual task switching, and visual and tactile working memory processing in congenitally deaf individuals.

An important unanswered question is whether modulation of crossmodal plasticity effects by task demands also occurs in older adults with ARHL. Theories of neuroplasticity propose that crossmodal reorganisation in sensory cortices primarily takes place during developmental ‘sensitive periods’ when plasticity is heightened, and that such plasticity declines with age (Knudsen, 2004; Burke & Barnes, 2006; Pascual-Leone et al., 2011; Hensch & Bilimoria, 2012). However, evidence from both animal and human studies demonstrates that crossmodal plasticity can also emerge following deafness or hearing loss in adulthood (Doucet et al., 2006; Allman et al., 2009; Lomber et al., 2010; Campbell & Sharma, 2014; Kim et al., 2016; Schormans et al., 2019). EEG studies have shown that visual and somatosensory stimuli evoke stronger responses in cortical auditory regions in older adults with hearing loss than in their full-hearing peers (Cardon & Sharma, 2018; Glick & Sharma, 2020). These studies have focused on basic sensory processing, and whether high-executive-demand plasticity effects occur outside sensitive developmental periods remains unknown. Understanding reorganisation in older adults with ARHL is particularly important given the strong association between ARHL and dementia. As such, it is crucial to determine not only how ARHL shapes crossmodal responses in auditory cortices, but also how crossmodal plasticity influences broader cognitive functions.

The aim of this study is to investigate whether there are crossmodal plasticity effects in auditory areas in older adults with ARHL, and whether this effect is modulated by task demands. We conducted an fMRI experiment in which older adults with and without ARHL performed a visual switching task and a visual working memory task. We hypothesised greater activity in auditory areas during conditions with higher executive demands in individuals with ARHL. Understanding plasticity and its modulation by task demands in the context of ARHL will inform the timescale of crossmodal reorganisation in the adult brain, determining whether these changes can occur outside developmental sensitive periods. It would also lay the groundwork for future investigations into the functional consequences of this reorganisation, which would determine whether crossmodal recruitment supports cognitive processing in other modalities, or whether it imposes additional demands on ageing neural systems. Clarifying these would provide a broader understanding of the cognitive impact of ARHL in ageing populations.

## Methods

All procedures followed standards set by the Declaration of Helsinki. Ethical approval was granted by the UCL Research Ethics Committee.

### Participants

There were two groups of participants (Supplementary Table 1):

a. Control group: participants with no ARHL (N=19; age: mean=70.4, range=65-81, SEM=1.15; 9 female, 10 male).
b. ARHL group: participants with hearing loss diagnosed after the age of 50 years old (N=18; age: mean=73.7, range=64-81, SEM=0.94; 11 female, 7 male). Hearing loss was defined according to the following criteria (Cruickshanks et al., 2003; Stevens et al., 2013): a) an average hearing threshold of more than 25 dB across 250, 500, and 1000 Hz; and b) an average hearing threshold above 35 dB across 250, 500, 1000, 2000, and 4000 Hz.

All participants were self-reported right-handed, native English speakers, with no history of psychiatric or neurological disorders, developmental language disorders or learning disabilities (N=37; age: mean=72.0, range=64-81, SEM=0.766; 20 female, 17 male). An independent samples t-test showed that the ARHL group is significantly older than the control group (p = 0.014; Supplementary Table 1). For this reason, age is added as a covariate in the behavioural and neuroimaging analysis.

Participants were recruited through adverts and invitation emails to relevant 3^rd^ sector organisations, and through the UCL Psychology subject pool. Prospective participants were sent a copy of the information sheet, a pre-screening questionnaire, and the MRI Safety Checklist. They were then contacted via telephone or video call to confirm their details and eligibility.

### Tasks

Participants performed two visual tasks that tap into two main components of executive function: working memory and task switching. Both tasks had one condition with higher executive demands (HED) and one with lower executive demands (LED).

The visual working memory task was a delayed-match-to-sample paradigm adapted from Manini et al. 2025. It was coded and presented using PsychToolBox in MATLAB 2023b (Brainard, 1997; Kleiner et al., 2007). The visual stimuli consisted of four 3.3° diameter circular gratings displayed on a grey background at ∼ 6.5° eccentricity (Figure 1). The gratings had a 30° tilt and were masked by a Gaussian mask. The texture of the gratings moved following a lateral uniform linear motion pattern, with the direction of motion changing at a frequency of 0.75-4.7 Hz. While the texture moved, the contour of the grating remained stationary, keeping the location of the whole grating constant.

**Figure 1.**
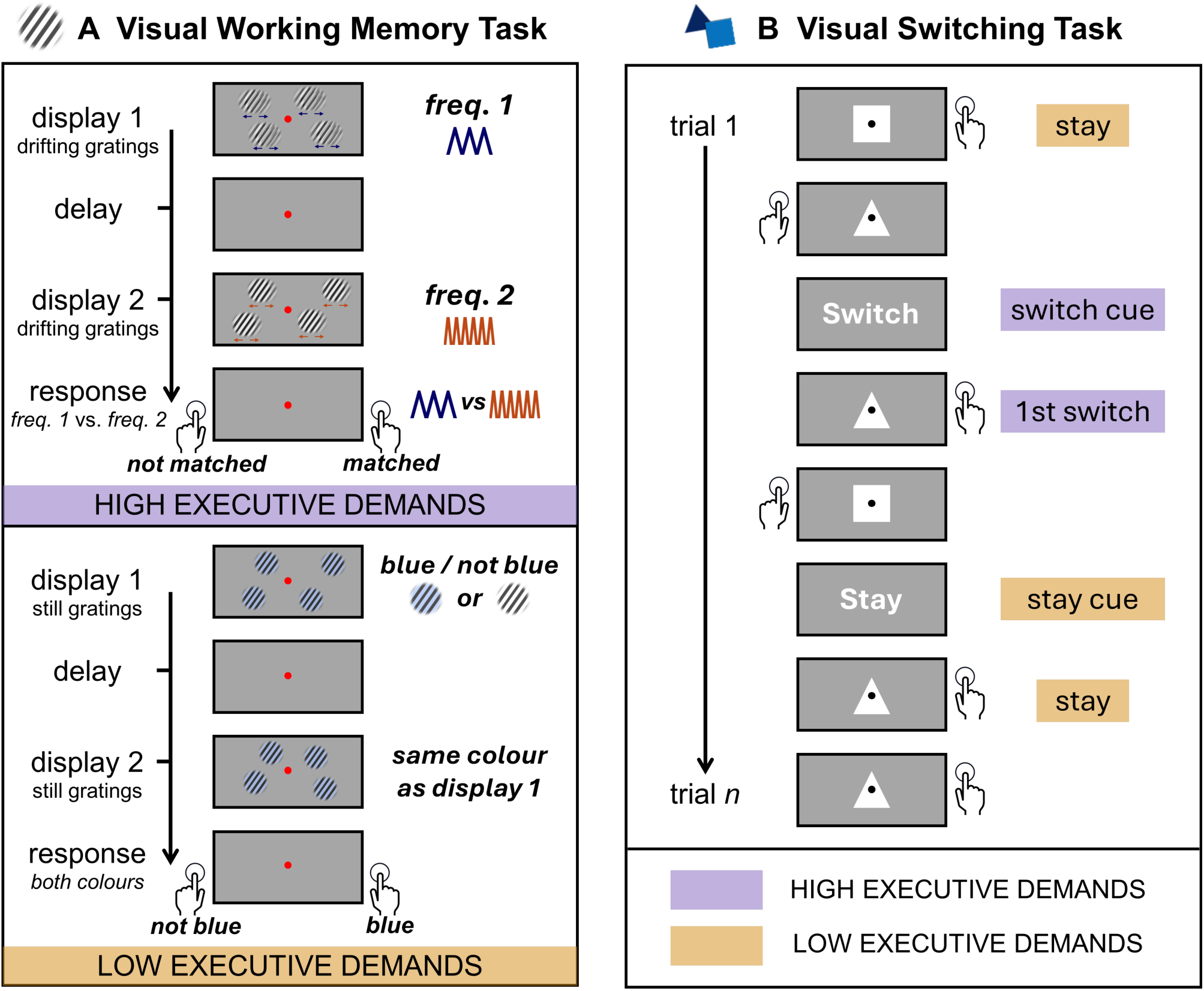
Experimental tasks. Diagrams of the working memory (A) and switching (B) tasks. Each task had a high executive demands condition (HED, purple) and a lower executive demands condition (LED, orange). See the ‘Methods’ section for details of the design.

During each trial, two visual stimuli were presented for 1000ms each, separated by a variable interstimulus interval (ISI) of 3000-5000ms (Figure 1). In the HED condition, participants had to remember the frequency of change (FoC) – that is the frequency at which the texture of the gratings changed its lateral direction of motion. At the end of each trial, participants had to indicate whether the 1^st^ stimulus (sample) and the 2^nd^ stimulus (probe) matched in their FoC. Samples had a FoC of either 1.5 or 3Hz. Probes FoC were: 0.5, 1.5, 3 or 4.7Hz. 50% of the trials had matching FoCs, and 50% had mismatching FoCs. In the LED condition, the texture of the gratings remained still, and in some trials the gratings were blue. At the end of each trial, participants had to indicate the colour of the gratings.

In both conditions, the gratings were displayed at variable polar angles; participants were told to fixate centrally and ignore the spatial location of the gratings. Participants responded with a right-handed button press, and the fixation dot turned from red to white to indicate that a response was recorded. Participants had up to 2500ms to enter their response, after which there was a jittered intertrial interval (ITI) of 1000-1400ms.

The visual switching task was adapted from previous studies (Rubinstein et al., 2001; Rushworth et al., 2002; Manini et al., 2022) and presented using PsychoPy 2023.2.3 (Peirce et al., 2019). On each trial, participants categorised a geometric shape (triangle or square; 1000 ms) with a corresponding button press within a 5000 ms response window. Visual feedback (500 ms) followed each response. Every 3-6 trials, a “Switch” or “Stay” cue (1000 ms) initiated a new block. “Switch” cues instructed participants to reverse the stimulus-response rule from the previous trial (HED), while “Stay” cues instructed them to maintain the current rule (LED). Inter-trial intervals jittered between 1000-2000 ms, and inter-block intervals jittered between 1000-2000 ms, interspersed with three randomly placed 6000-ms intervals. Each run consisted of 160 trials, divided into 20 switch (HED) and 20 stay (LED) blocks in random order.

### Experiment procedure

Each experimental session included the following behavioural tests:

i Task familiarisation: Participants learned and practised each task through a computer-based interactive tutorial demo, followed by a practice run. They were given additional verbal or written instructions, if required. All participants fully understood the task and reached at 80% of accuracy in their final practice run.
ii Non-verbal IQ test: This was evaluated using Matrix Reasoning subtest of the Wechsler Abbreviated Scale of Intelligence (WASI, Wechsler, 1999). There was no significant difference between groups in their converted WASI scores (*t* = 0.654, *p* = 0.517).
iii Pure tone audiogram (PTA): PTAs were conducted using a Resonance R17 screening portable audiometer following the Recommended Procedure of the British Society of Audiology (British Society of Audiology, 2018). All participants were tested for each ear unaided in six frequency bands (250 Hz, 500 Hz, 1000 Hz, 2000 Hz, 4000 Hz, 8000 Hz).
iv Speech-in-noise test (SIN): Sentence recognition was tested using Institute of Electrical and Electronics Engineers (IEEE) sentences (Institute of Electrical and Electronics Engineers, 1969) recorded from a male talker of Standard Southern British English presented against a background of 20-talker babble. Tests were conducted using custom programmes on MATLAB 2023a. Sounds were presented via Epos Sennheiser Impact SC 665 headset. On each trial, the masker commenced 700 ms before the onset of the target speech and continued for 200 ms after the target offset, with cosine onset and offset ramps of 100 ms applied to the mixture. Participants’ spoken responses were scored by an experimenter. Each sentence contained five key words on which scoring was based. An adaptive procedure was used to estimate the Sentence Reception Threshold (SRT), defined as the signal-to-noise ratio (SNR) at which 50% of key words were correctly identified. The first sentence in each measurement run was presented at an SNR of + 10 dB. Thereafter, the SNR decreased if more than half of the keywords were correctly identified and increased otherwise. A 10-dB change in SNR was used until the first reversal, 6.5 dB until the second reversal, and 3 dB for all subsequent reversals. The procedure terminated after 8 reversals with the 3-dB step size or after a maximum of 20 sentences. SRTs were calculated as the mean SNR for the final even number of reversals with the 3-dB step size.

**Figure 2.**
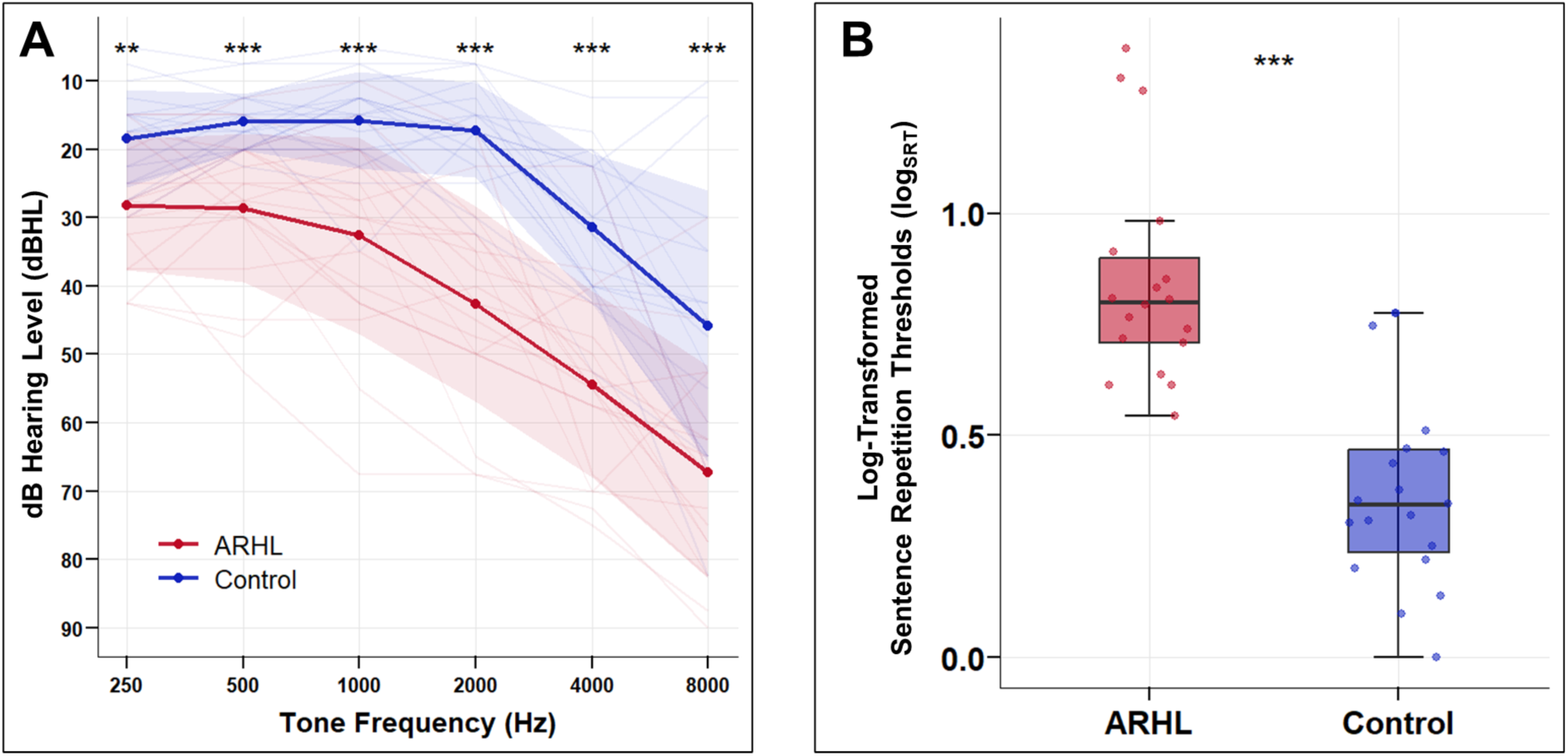
Pure-Tone Average (PTA) and Log-Transformed Sentence Repetition Thresholds (Log_SRT_). **A.** The bold lines show the average pure-tone hearing level for the ARHL group (red) and control group (blue) for of both ears. Thin lines show data from each individual participant. **B.** Log-transformed speech in noise thresholds. On each box, the edges indicate the 25^th^ and 75^th^ percentiles, and the black horizontal line inside the box indicates the median. Asterisk signs for student’s t-test results between groups (hypothesis: *µ*_*ARHL*_ ≠ *µ*_*Control*_): ***p<.001 **p<.01.

Scanning took place at Birkbeck - UCL Centre for Neuroimaging (BUCNI). MRI data were obtained on a 1.5 T Siemens Avanto scanner with a 32-channel head coil. Researchers communicated with participants in the scanner via an intercom or via written text on a screen; participants responded verbally. All participants were given ear protectors.

Each scanning session included 1-2 runs of each task, one anatomical scan, one sagittal localiser and one breath-hold scan (not reported here). Functional imaging data during the tasks were acquired using a two-dimensional echo planar imaging sequence [time to repeat (TR): 1000 ms, time to echo (TE): 54.8 ms, flip angle (FA): 75 deg, field of view (FoV): 205 × 205 mm^2^, 3.2 × 3.2 × 3.2 mm^3^ resolution, 40 slices]. Structural imaging data were acquired via magnetisation-prepared rapid acquisition with gradient echo (MPRAGE, TR: 2730 ms, TE: 3.57 ms, FA: 7 deg, FoV: 224 × 256 mm^2^, 1 × 1 × 1 mm^3^ resolution, 176 slices).

### Statistical analysis of behavioural results

Average accuracy and reaction time were compared between groups (Control and ARHL) and conditions (HED and LED).

Behavioural performance in the working memory and switching tasks was analysed using repeated-measures ANOVAs. For the working memory task, accuracy served as the dependent variable, whereas for the switching task, both accuracy and reaction time were analysed separately as dependent variables. Missed trials and trials with response times longer than 2000 ms or shorter than 200 ms were excluded from the analyses. Condition (HED and LED) was entered as a within-subject factor and Group (Control and ARHL) as a between-subjects factor. Tukey’s post hoc tests were conducted for all ANOVAs, and age was included as a covariate.

All statistical analyses were performed using manual scripts or default built-in functions on MATLAB 2023a, R 4.4.0 via RStudio (Posit team, 2024; R Core Team, 2024), and Jamovi-2.3.28.0 (The Jamovi Project, 2022).

### MRI preprocessing

All MRI data were analysed using MATLAB 2023a and SPM12 (Penny et al., 2011).

For each participant’s anatomical scan, segmentation and normalisation were conducted with SPM12’s standard procedures. Skull-stripping was done via the Image Calculation function (ImCalc, http://tools.robjellis.net). The expression we used was:

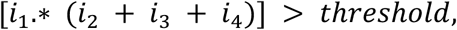

in which *i*_1_ was the bias-corrected structural scan while *i*_2_, *i*_3_ and *i*_4_ represented the tissue images (grey matter, white matter and cerebrospinal fluid).

The functional data were realigned and coregistered via SPM12’s standard procedures and then referred and normalised to the standard Montreal Neurological Institute (MNI) space. To allow time for the magnetisation to reach a dynamic equilibrium, the first seven images of each run were manually excluded.

### Functional MRI analysis

All model specifications and estimation were conducted using SPM12. A first-level analysis was conducted for each participant and each task by fitting a general linear model (GLM) with regressors for each condition of interest. For both tasks, onsets of all HED and LED trials were entered into the GLM as regressors of interest. The corresponding events for each regressor were modelled as a boxcar of the relevant duration and convolved with the canonical haemodynamic response function in SPM. Motion parameters were entered as regressors of no interest. Multiple regression was used to generate parameter estimates for each regressor at every voxel. For each task, subject-specific images generated in the first-level analysis were entered into a full factorial second-level analysis.

Visualisation and rendering were done via the bspmview toolbox (Spunt, 2016). We applied specified t-contrasts to assess the effects and interactions of interest. Coordinates of the peaks of activation are reported in the form of MNI x, y and z coordinates. Regions were automatically labelled using the automated anatomical atlas 3 (AAL3, Rolls et al., 2020).

### Region of interest analysis

Bilateral Te1.0 definitions were obtained from the Julich-Brain Cytoarchitectonic Atlas 3.1 (Amunts et al., 2020, 2023), in which Te1.0 was distinguished from other auditory areas based on cytoarchitectonic characteristics (Zachlod et al., 2020).

Bilateral pSTC was defined based on the findings of Cardin et al., 2018, where a visual working memory crossmodal plasticity effect was found in bilateral pSTC in early deaf individuals. Data from Cardin et al., 2018 was used to define right and left pSTC ROIs with the contrast [deaf (working memory > control task) > control (working memory > control task)] (p < 0.005, uncorrected).

Values from each contrast of interest were extracted for each participant and each ROI using MarsBaR v.0.45 toolbox (http://marsbar.sourceforge.net) (Brett et al., 2002).

Repeated measures ANOVAs were used to test statistical significance of the effects. To test the effects of group, in the first set of ANOVAs, condition (HED, LED) was entered as repeated measures factors; group (ARHL, Control) was entered as a between-subject factor; age was entered as a non-interactive covariate. To investigate the relationship between specific hearing measurements and the BOLD response in auditory regions, we conducted a second set of ANOVAs, where condition was entered as repeated measures factor and PTA or SRTs, and age, were entered as non-interactive covariates.

All statistical analyses were performed using manual scripts or default built-in functions on MATLAB 2022a, R 4.4.0 via RStudio, and Jamovi-2.3.28.0.

## Results

### Behavioural results

Behavioural results from the participants’ performance in the scanner are shown in Figure 4. To examine differences between groups and conditions, we performed separate repeated measures ANOVAs, with either accuracy (for each task) or reaction time (for switching only) as the dependent variable (Supplementary Table 2). Group (ARHL and Control) was entered as a between-subject factor, Condition (HED and LED) was entered as a within-subject factor, and age was entered as a non-interactive covariate. Results show a significant main effect of condition for accuracy in both tasks [*Working Memory*: *F*(1,31) = 42.87, *p* < 0.001, 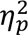 = 0.580; *Switching*: *F*(1,34) = 27.97, *p* < 0.001, 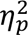 = 0.451], and for reaction time in the switching task [*F*(1,35) = 67.13, *p* < 0.001, 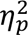 = 0.664]. These effects confirm the higher cognitive demands of the HED conditions. No significant effect of group was observed.

**Figure 3.**
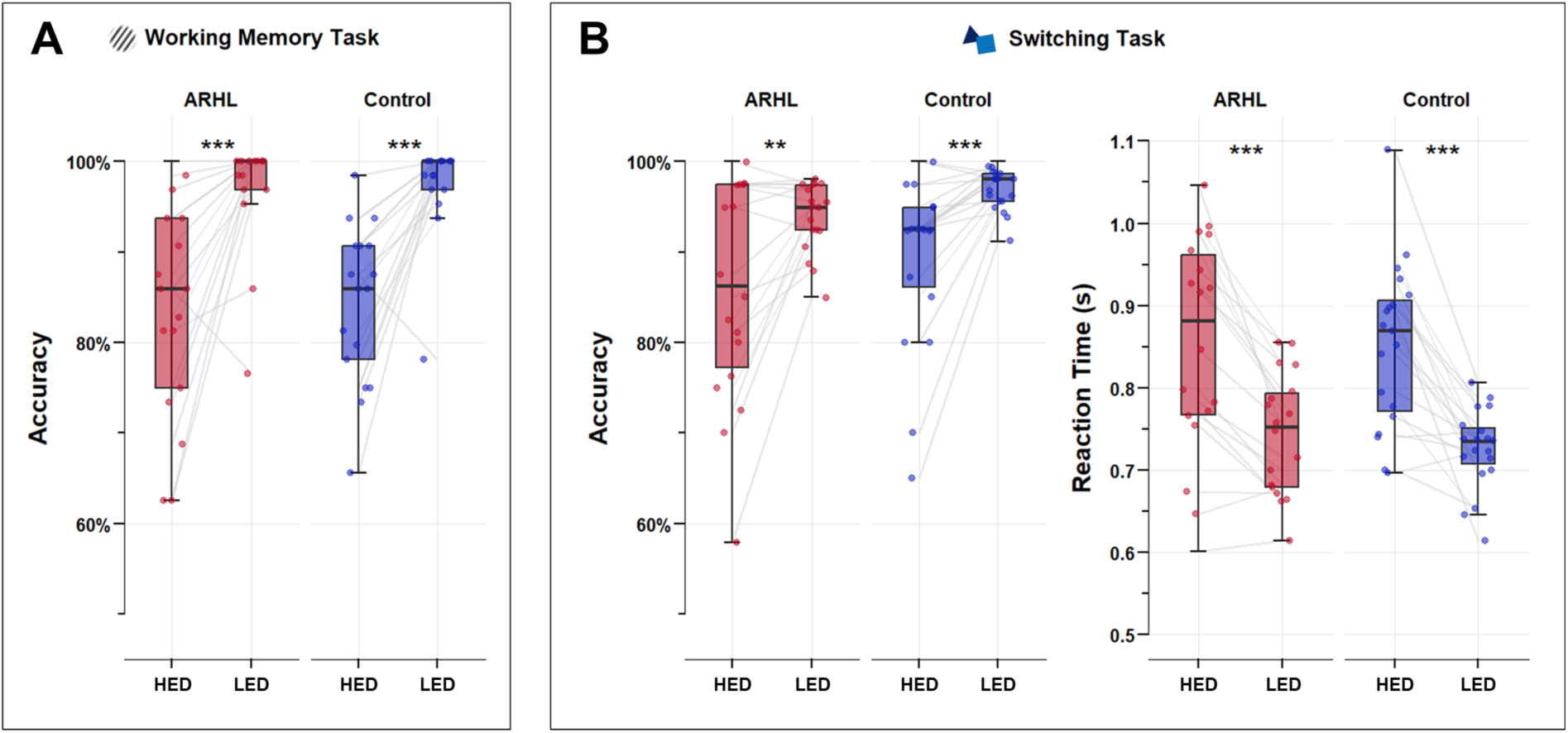
Behavioural Performance. The data points show an individual’s average performance in each condition. On each box, the edges indicate the 25^th^ and 75^th^ percentiles, and the black horizontal line inside the box indicates the median. ARHL: Age-related hearing loss. HED: high executive demands; LED: low executive demands. Asterisk signs for Tukey’s Honestly Significant Difference test results between conditions (hypothesis: *µ*_*HED*_ ≠ *µ*_*LED*_): ***p<.001 **p<.01.

**Figure 4.**
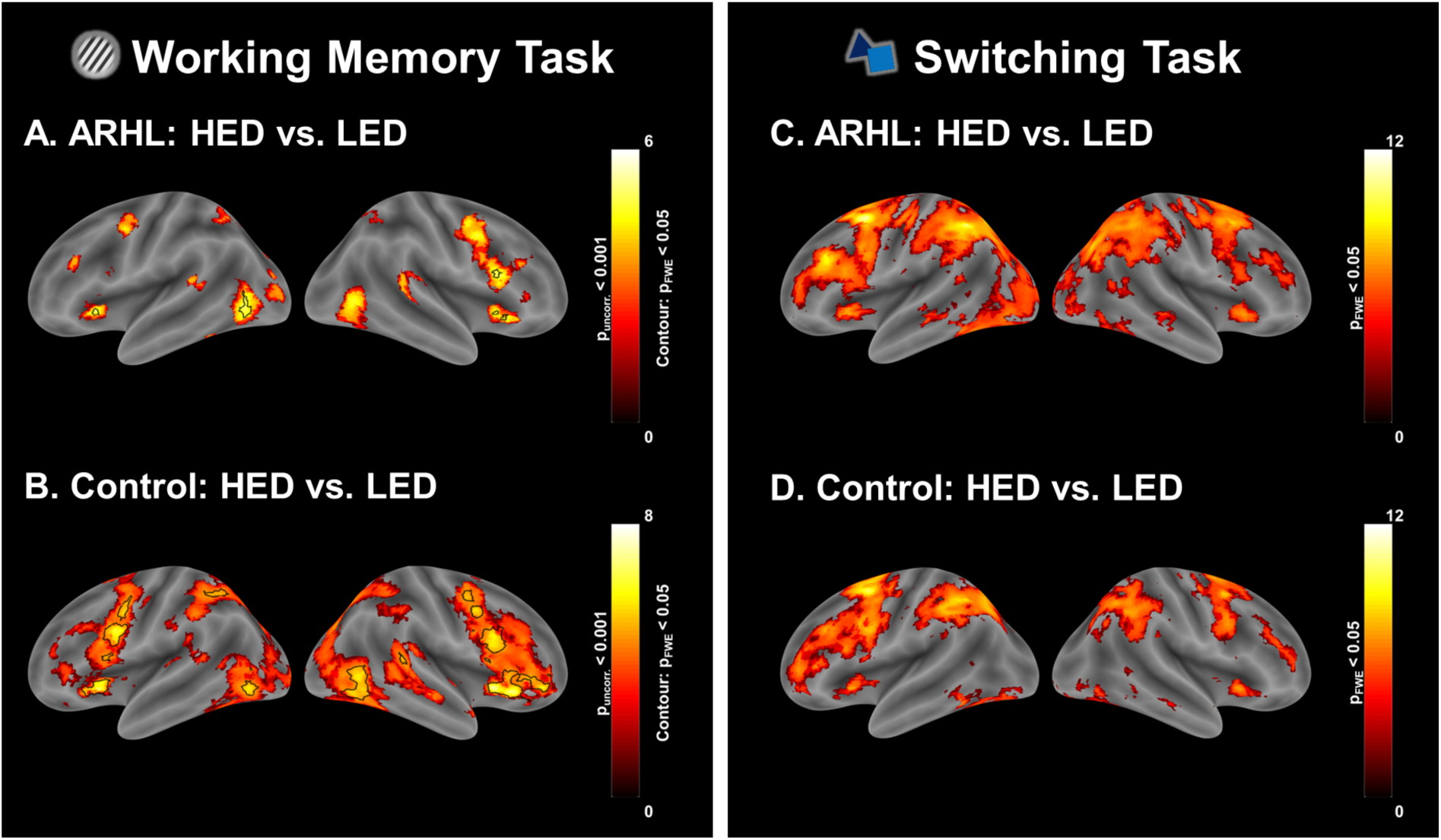
Whole brain results. The figure shows whole-brain activations for the contrast [HED > LED] for each group and each task. Left Panel: Results from the contrast [Working Memory (HED) > Control (LED)]. Results are shown at p<.001 for display purposes, but all peaks were significant at p<.05 (FWE corrected; black contour). Right Panel: Results from the contrast [Switch (HED) > Stay (LED)]. Results are shown on at p<.05 FWE corrected. Colour bars represent z-scores. HED: high executive demands; LED: low executive demands.

### Functional MRI results – Whole Brain Analysis

In both groups combined, the contrasts [HED vs LED] in the switching and in the working memory tasks elicited activation in frontoparietal regions typically involved in executive functions, as observed in previous studies (Figure 4; Braver et al., 2003; Gold et al., 2010; Manini et al., 2022, 2025).

Group comparisons in the switching and working memory tasks for the contrasts [HED vs LED] revealed no significant differences between groups (FWE > .05).

### Functional MRI results – ROI analysis

To further explore differences between conditions and groups in auditory areas, we conducted a ROI analysis on bilateral Te1.0 and bilateral pSTC for both tasks. Contrasts values for each ROI were entered into a repeated measures ANOVAs with Group (ARHL and Control) as a between-subject factor, Condition (HED and LED) as a within-subject factor and age as a non-interactive covariate.

For the working memory task (Supplementary Table 3), bilateral Te1.0 showed no effect of condition [*Left Te*1.0: *F*(1,31) = 4.13, *p* = 0.051; *Right Te*1.0: *F*(1,31) = 0.242, *p* = 0.626], while bilateral pSTC showed a significant effect of condition [*Left pSTC*: *F*(1,31) = 18.46, *p* < 0.001, 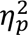 = 0.373; *Right pSTC*: *F*(1,31) = 17.78, *p* < 0.001, 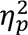 = 0.364]. No significant effect of group or age was observed.

For the switching task (Supplementary Table 3), results from all four ROIs showed a significant effect of condition [*Left Te*1.0: *F*(1,34) = 17.36, *p* < 0.001, 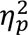 = 0.338; *Right Te*1.0: *F*(1,34) = 10.38, *p* < 0.001, 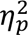 = 0.234; *Left pSTC*: *F*(1,34) = 20.31, *p* < 0.001, 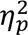 = 0.374; *Right pSTC*: *F*(1,34) = 22.23, *p* < 0.001, 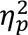 = 0.395]. No significant effect of group or age was observed.

One-sample t-tests revealed that in the switching task, the HED condition was significantly different from zero in all ROIs (Figure 5 C & D). However, for the WM task, these differences were significant only left Te1.0 and left pSTC (Figure 5 A & B).

**Figure 5.**
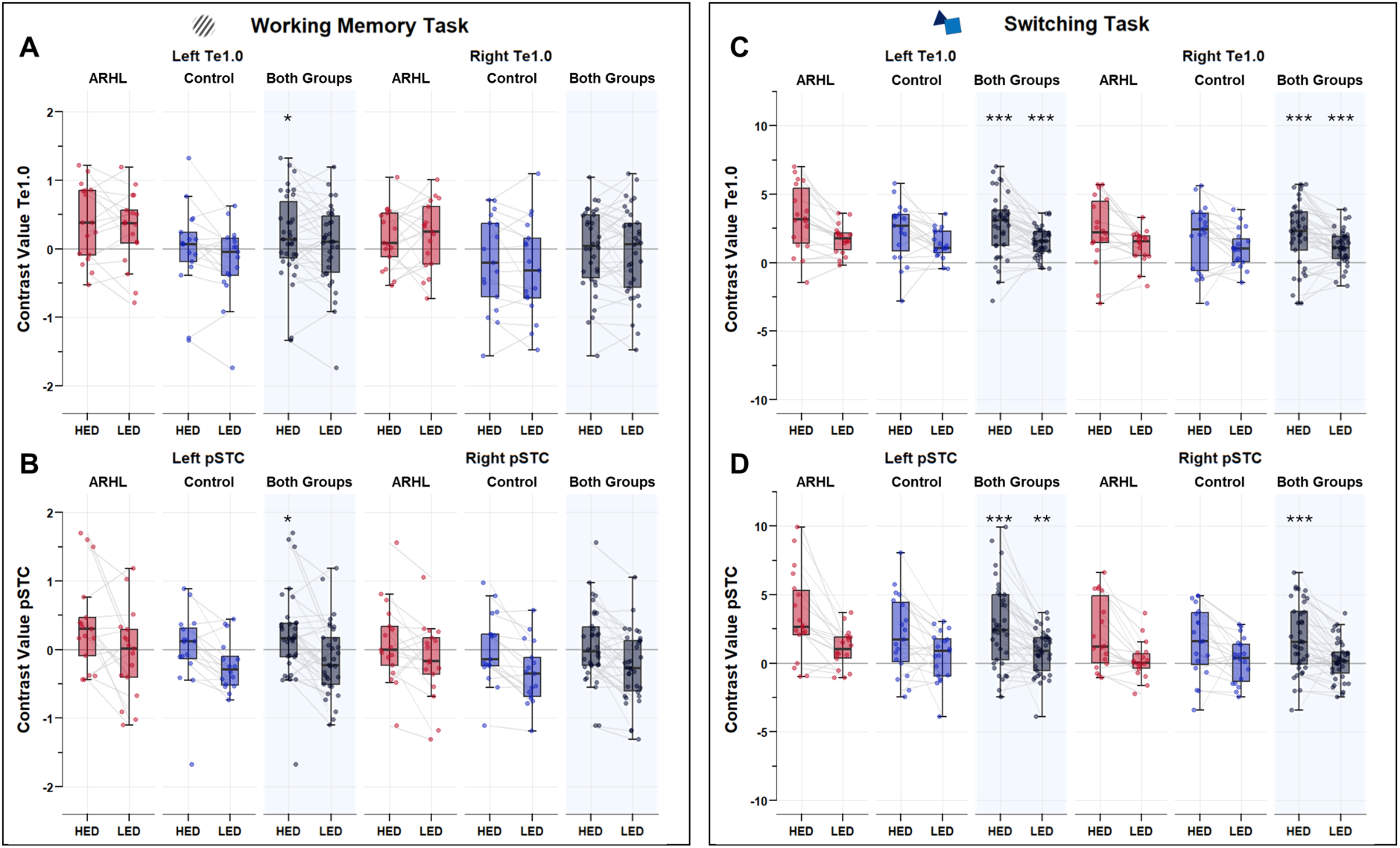
Region of Interest (ROI) Analysis. The data points show an individual’s average contrast values for a given condition vs the implicit baseline. On each box, the edges indicate the 25^th^ and 75^th^ percentiles, and the black horizontal line inside the box indicates the median. Notice the different y-axes in the working memory and switching tasks. ARHL: Age-related hearing loss. HED: high executive demands; LED: low executive demands. pSTC: posterior superior temporal cortex. Asterisk signs for one-sample t-test results (hypothesis: *µ > 0)*: ***p<.001, **p<.01, *p<.05.

To further explore the effect of hearing loss on crossmodal plasticity effects in auditory areas, we conducted a second set of repeated measures ANOVAs where condition was entered as repeated measures factor and PTA threshold or SRTs were entered as covariates (the between-subject factor Group was removed). Age was entered as a covariate. There was no significant effect of SRT or PTA in any of the ROIs (p > .05; Supplementary Table 4).

## Discussion

In this study, we examined crossmodal plasticity effects in auditory cortical regions of older adults with age-related hearing loss (ARHL). Specifically, we tested whether auditory areas are recruited during two visual executive function tasks, each comprising a high executive demands condition (HED) and a low executive demands condition (LED). As expected, HED conditions elicited stronger activations in frontoparietal brain areas typically involved in executive processing and cognitive control. Our results also show that the HED condition activated cortical areas typically involved in auditory processing in both groups of participants. Specifically, the HED condition of the switching task and the working memory tasks elicited increased activation in bilateral pSTC. In Te1.0, only the switching task showed a stronger activation in the HED task. While previous studies have suggested the presence of crossmodal plasticity in older adults, our results demonstrate that this is modulated by task demands, where activations are more significant for higher-order cognitive functions. We showed anatomical specificity in the effects, showing that high-executive-demand crossmodal effects in older adults are present in primary and secondary auditory areas in the switching task, but only found in pSTC in the working memory task.

Overall, these findings show high-executive-demand crossmodal plasticity effects, similar to those found in congenitally deaf individuals, also in the auditory cortex of older adults with and without hearing loss. Previous studies of younger adults without hearing loss did not reveal such a crossmodal effect in the age-matched control group (Manini et al. 2022, Manini et al. 2025). As such, results from this study suggest that the observed crossmodal recruitment of auditory regions can be due either to: 1) effects of age-typical hearing loss that were present in both groups; 2) a general effect of ageing, or 3) a combination of the two. In the following sections, we discuss these possibilities in detail.

### Effects of age-related hearing loss in the crossmodal reorganisation of the auditory cortex

Crossmodal plasticity is well documented in congenitally deaf individuals, including recruitment of auditory regions during high-demand visual and somatosensory tasks (Ding et al., 2015; Cardin et al., 2018; Manini et al., 2022; Manini et al., 2025). Crossmodal effects are commonly found during ‘sensitive periods’ of enhanced plasticity in early life (Hensch, 2004; Knudsen, 2004; Hensch & Bilimoria, 2012). Here we investigated whether similar reorganisation can also emerge later in life (after the critical periods of sensory development have passed), and whether this reorganisation is modulated by task demands. Our findings suggest that this may be the case: older adults with ARHL showed increased activation in auditory cortical regions during visual executive function tasks. Although comparable effects were observed in the control group, this does not rule out hearing loss as a key contributor. Most older adults with age-typical hearing present a degree of high-frequency hearing loss (Humes et al., 2010; Humes, 2019). This is also evident in our sample, where the average PTA threshold at 4kHz and 8kHz is above 25 dBHL. A possible explanation for crossmodal plasticity effects is that the degree of hearing loss observed in older adults who are diagnosed with ARHL may be sufficient to drive plasticity in the auditory cortex. Supporting this interpretation, Glick and Sharma (2020) reported significant differences between individuals with and without hearing loss in the peak latency of the ERP P1 component evoked by visual motion, with the latency negatively correlated with high-frequency PTA. In their study, the control group did not exhibit high-frequency hearing loss (but notice that the average age of their cohort was eight years younger – mean = 64 years SD = 4.68, compared to mean = 72.0 years, SD = 4.72). These findings suggest that the mild high-frequency hearing loss present in our control group could be sufficient to induce visual crossmodal responses in auditory regions.

In congenitally deaf individuals, a range of potential mechanisms has been discussed to explain high-executive demand crossmodal plasticity effects (see Cardin for a review). These include top-down influences from frontoparietal regions (Cardin et al., 2023; Friederici, 2011; Skeide & Friederici, 2016), the anatomical proximity between auditory cortices and temporoparietal regions involved in attentional reorienting and task switching (Cardin et al., 2020; Patel et al., 2019; Shiell et al., 2016; Vinogradova & Cardin, 2024), and the preservation of early-developing functions following reduced functional differentiation for auditory processing (Cardin et al., 2020). While such crossmodal effects are especially pronounced during developmental “sensitive periods” (Knudsen, 2004), an important question is how hearing loss can still drive crossmodal reorganisation in older adults, long after these critical periods have passed. One mechanism that may account for this later-life plasticity is hyperactivity and reduced inhibition in the auditory pathway (Herrmann & Butler, 2021). In hearing loss, the reduction in auditory inputs results in increased neural excitability in the auditory cortex (Herrmann & Butler, 2021). This hyperactivity is thought to be a compensatory mechanism, whereby the brain attempts to enhance sensitivity to available sensory inputs by increasing activity along the auditory pathway. Research shows that hyperactivity in the central auditory system is associated with reduced synaptic inhibition (Auerbach et al. 2014; Ouda et al. 2015; Sanes et al. 2010; Salvi et al. 2017). This reduced inhibition somehow resembles the neurophysiological environment of early development when inhibitory circuits are still maturing. Because reduced inhibition is strongly associated with increased plasticity, hearing loss in older adults may create a cortical environment conducive to reorganisation, enabling auditory regions to engage in crossmodal cognitive processes. This could be achieved by amplifying preexisting higher-order representations already present in the auditory cortex. In previous work (Manini et al., 2025), we showed that not only congenitally deaf individuals but also hearing individuals exhibit abstract, non-feature-specific representations of crossmodal task information in auditory areas. While the functional relevance of this crossmodal information seems to be enhanced in congenitally deaf individuals, increased cortical excitability due to hearing loss could also enhance the functional relevance of such latent representations in older adults with hearing loss, contributing to the recruitment of auditory regions during executive tasks.

### Effect of ageing on crossmodal plasticity in the auditory cortex

We found a significant effect of condition in the recruitment of auditory regions pSTC and Te1.0 during the switching task in both the control and ARHL groups. In both these areas, activations in the HED condition were significantly different from baseline. In the working memory task, there was a significant effect of condition in bilateral pSTC, but only in left pSTC this was accompanied by a significant difference from baseline. In our previous studies examining crossmodal plasticity during a switching task, auditory regions were activated in congenitally deaf adults, but not in age-matched hearing control groups (Manini et al., 2022, 2025). A possible explanation for the findings in the control group in this study is that crossmodal plasticity effects are due to age-related changes in the auditory cortex of older adults, independently of hearing loss.

Cognitive neuroscience studies of human ageing can provide a conceptual framework for our findings. The observed increase in activation in auditory areas during visual cognitive tasks could reflect compensatory mechanisms usually observed in older adults during high-demand cognitive tasks (Qin & Basak, 2020). Previous studies have shown increased activations across a wider network of brain regions in the older adult brain during cognitive tasks (Qin & Basak, 2020; McClaskey, 2024). The potential relevance of these ‘compensatory’ recruitment is to sustain performance to a level comparable of that of younger adults. Under this framework, auditory areas may be recruited to support performance. In addition, studies of ageing have shown less differentiated neural responses to distinct stimulus categories (dedifferentiation hypothesis; see Koen & Rugg, 2019 for a review). While this has been mostly documented in the visual domain and in the study of object categories, a similar mechanism can be conceived at a crossmodal level, where the sensory cortices show ‘less differentiated’ response for their main sensory modality.

At a physiological level, ageing, independently of hearing loss, is also associated with disinhibition in the auditory cortex (Ibrahim & Llano, 2019). As explained in the section above, disinhibition can enhance plastic reorganisation in the auditory cortex. A reduction in GABAergic synaptic inhibition has been observed in the auditory cortex in animal models of ageing (Ouda et al., 2015; Ouellet & de Villers-Sidani, 2014; Brewton et al., 2016). This reduction could be due to a gradual decline of the cellular energy metabolism as individuals age. Inhibitory neurons are known to be metabolically demanding, and particularly vulnerable to age-related metabolic decline, resulting in reduced inhibitory control and heightened spontaneous activity (Ibrahim & Llano, 2019). This hyperexcitability could resemble the neural state characteristic of developmental periods in which sensory cortices are highly plastic. Consequently, older adults may exhibit a renewed capacity for functional reorganisation, allowing auditory regions to respond to non-auditory cognitive demands. This mechanism could provide an explanation how auditory hyperactivity emerges with ageing, even in the absence of peripheral hearing loss (Herrmann & Butler, 2021; Rogalla & Hildebrandt, 2020). In summary, ageing can not only create additional computational demands, but also the metabolic and physiological environment for crossmodal plasticity effects to arise.

### Combined effects of ageing and age-related hearing loss

While we have discussed the effects of hearing loss and ageing separately, in practice, chronological age and hearing loss are highly correlated. This tight relationship makes it challenging to isolate the independent effects of hearing loss and chronological age on the function of the auditory cortex in older adults. A further complication is that many studies of ageing do not assess or control for hearing ability, meaning that findings attributed to “ageing” may, in fact, reflect unmeasured hearing loss (see Cardin, 2016, for discussion). As such, the conceptual distinction between ageing and hearing loss becomes blurred, because hearing loss is itself a major physiological change that defines ageing. Considering the above, it is likely that the observed crossmodal activity in auditory regions in older adults is due to the combined effects of reduced auditory inputs and ageing effects. Hearing loss could reduce the sensory constraints on the auditory cortex, while ageing could broaden the reliance on distributed cortical networks during cognitive tasks. Together, these factors create a scenario in which auditory regions become more available and likely to participate in high-demand processing across modalities.

### Switching preferentially drives crossmodal effects in older adults

While previous studies have suggested crossmodal plasticity in auditory areas in older adults (Cardon & Sharma, 2018; Glick & Sharma, 2020), our results demonstrate this effect specifically for higher-order cognitive functions. The differences we observed between the switching and working memory tasks align with our prior findings in congenitally deaf individuals, where we have shown that not all executive function tasks activate auditory areas in deaf individuals, and that switching tasks are particularly effective at revealing crossmodal effects (Manini et al., 2022). In our previous research in congenitally deaf individuals, both conditions of the visual switching task (Switch and Stay) activated auditory regions, including pSTC and primary auditory cortex, with significantly stronger activation during the Switch condition (Manini et al., 2022). Similarly, in the present study, the switching task produced more robust crossmodal responses in auditory cortex than the visual working memory task. This may be due to the fast temporal updating nature of the stimuli in the task, which may tap into the specialisation of the auditory cortex for temporal processing, or into a crossmodal role of auditory regions in updating and reallocation of attention (see Manini et al., 2022, for a discussion). Indeed, even with the delay-to-match working memory task, a temporal version of the task drives the auditory cortex more strongly than a spatial version of the task that contains the same sensory features (Manini et al., 2025). This may explain why previous studies of visual working memory in older adults with hearing loss (Rosemann & Thiel, 2020) reported no activations in auditory areas, possibly because of the use of a recognition-based task, which may now have the temporal dynamics and needed updating that could drive crossmodal activations in auditory areas. Together, these findings highlight that crossmodal plasticity in older adults is modulated by task demands, being stronger for tasks that require temporal updating and reallocation of attention, such as task switching.

## Conclusion

Our findings show that auditory cortical regions in older adults are engaged during high-demand visual tasks in individuals with and without ARHL. This indicates that high-executive demands crossmodal reorganisation is not confined to early developmental periods and may emerge in later life, when auditory input declines and executive demands increase.

Understanding crossmodal plasticity in the auditory cortex and its role in higher-order cognition is crucial not only for its potential effects on auditory processing but also given the established relationship between ARHL and dementia. This work provides a foundation for future research investigating whether crossmodal plasticity supports cognitive performance or whether it imposes additional processing demands. Overall, our findings highlight the importance of considering both sensory loss and ageing-related neural changes in brain function in older adults, improving our understanding of sensory and cognitive processing in ageing.

## Ethical statement

All participants provided written consent prior to participation and were compensated for their time. All experiments were conducted in accordance with the standards set forth by the Declaration of Helsinki and obtained approval from the UCL Psychology Ethics Steering Committee.

## Supplementary Materials

**Supplementary Table 1.** Demographics.

|  | Age |  | Gender | WASI |  |
| --- | --- | --- | --- | --- | --- |
|  | Mean (Range) | SD |  | Mean | SD |
| **Full Sample**<br>N = 37 | 72.0 (64-81) | 4.72 | 20F : 17M | 61.9 | 8.34 |
| **Switching**<br>ARHL N = 18 | 73.7 (64-81) | 3.97 | 11F : 7M | 62.8 | 8.15 |
| **Switching**<br>Control N = 19 | 70.4 (65-81) | 4.89 | 9F : 10M | 61.0 | 8.65 |
| **Working Memory**<br>ARHL N = 17 | 73.8 (64-81) | 4.09 | 11F : 6M | 63.1 | 8.35 |
| **Working Memory**<br>Control N = 17 | 70.0 (65-81) | 5.02 | 8F : 9M | 60.5 | 9.08 |

*Note.* Three participants (1 in ARHL, 2 in control) were excluded from working memory task analysis due to low accuracy (<60%).

Group Comparisons
|  | N |  | Age |  | Gender |  | WASI |  |
| --- | --- | --- | --- | --- | --- | --- | --- | --- |
|  | ARHL | Control | *t* | *p* | *X*<sup>2</sup> | *p* | *t* | *p* |
| **Working Memory** | 17 | 17 | **2.397** | **0.023** | 0.471 | 0.493 | 0.844 | 0.405 |
| **Switching** | 18 | 19 | **2.283** | **0.029** | 0.243 | 0.622 | 0.654 | 0.517 |

*Note.* Significant effects are shown in bold. One participant in the control group did not complete the WASI test.

**Supplementary Table 2.** Repeated Measures ANOVAs of Behavioural Results.

|  | Working Memory |  |  | Switching |  |  |  |  |  |
| --- | --- | --- | --- | --- | --- | --- | --- | --- | --- |
|  | Accuracy |  |  | Accuracy |  |  | Reaction Time |  |  |
| | *F* (*df*) | *p* | $\eta^2_p$ | *F* (*df*) | *p* | $\eta^2_p$ | *F* (*df*) | *p* | $\eta^2_p$ |
| Condition | **42.869** (1,31) | **< .001** | **0.580** | **27.973** (1,34) | **< .001** | **0.451** | **67.127** (1,34) | **< .001** | **0.664** |
| Group | 0.057 (1,31) | 0.813 | 0.002 | 1.772 (1,34) | 0.192 | 0.050 | 0.009 (1,34) | 0.923 | 0.000 |
| Age | 1.986 (1,31) | 0.169 | 0.060 | 0.030 (1,34) | 0.864 | 0.001 | 0.347 (1,34) | 0.560 | 0.010 |

*Note.* Type 2 Sums of Squares. Factors in the analysis were: condition (HED, LED), group (ARHL, control), and age. Significant effects are shown in bold.

**Supplementary Table 3.** Repeated Measures ANOVAs for ROI analysis.

| Working Memory |  |  |  |  |  |  |  |  |  |  |  |  |
| --- | --- | --- | --- | --- | --- | --- | --- | --- | --- | --- | --- | --- |
|  | Left Te1.0 |  |  | Right Te1.0 |  |  | Left pSTC |  |  | Right pSTC |  |  |
| | *F* (*df*) | *p* | $\eta^2_p$ | *F* (*df*) | *p* | $\eta^2_p$ | *F* (*df*) | *p* | $\eta^2_p$ | *F* (*df*) | *p* | $\eta^2_p$ |
| Condition | 4.133 (1,31) | 0.051 | 0.118 | 0.242 (1,31) | 0.626 | 0.008 | **18.459** (1,31) | **< .001** | **0.373** | **17.777** (1,31) | **< .001** | **0.364** |
| Group | 2.914 (1,31) | 0.098 | 0.086 | 2.380 (1,31) | 0.133 | 0.071 | 0.350 (1,31) | 0.559 | 0.011 | < .001 (1,31) | 0.986 | < .001 |
| Age | 0.923 (1,31) | 0.344 | 0.029 | 1.930 (1,31) | 0.175 | 0.059 | 2.256 (1,31) | 0.143 | 0.068 | 3.580 (1,31) | 0.068 | 0.104 |
| Switching |  |  |  |  |  |  |  |  |  |  |  |  |
|  | Left Te1.0 |  |  | Right Te1.0 |  |  | Left pSTC |  |  | Right pSTC |  |  |
| | *F* (*df*) | *p* | $\eta^2_p$ | *F* (*df*) | *p* | $\eta^2_p$ | *F* (*df*) | *p* | $\eta^2_p$ | *F* (*df*) | *p* | $\eta^2_p$ |
| Condition | **17.360** (1,34) | **< .001** | **0.338** | **10.378** (1,34) | **0.003** | **0.234** | **20.311** (1,34) | **< .001** | **0.374** | **22.230** (1,34) | **< .001** | **0.395** |
| Group | 2.510 (1,34) | 0.123 | 0.069 | 0.613 (1,34) | 0.439 | 0.018 | 1.213 (1,34) | 0.279 | 0.034 | 0.174 (1,34) | 0.679 | 0.005 |
| Age | 1.070 (1,34) | 0.308 | 0.031 | 0.558 (1,34) | 0.460 | 0.016 | 0.255 (1,34) | 0.617 | 0.007 | 0.006 (1,34) | 0.941 | 0.000 |

Note. Type 2 Sums of Squares. Factors in the analysis were: condition (HED, LED), group (ARHL, control), and age. Significant effects are shown in bold.

**Supplementary Table 4.** Repeated Measures ANOVAs for ROI analysis with PTA or SRT as covariates.

|  | Working Memory |  |  |  | Switching |  |  |  |
| --- | --- | --- | --- | --- | --- | --- | --- | --- |
|  | PTA |  | SRT |  | PTA |  | SRT |  |
|  | *F (df)* | *p* | *F (df)* | *p* | *F (df)* | *p* | *F (df)* | *p* |
| Left Te1.0 | 0.239 (1,31) | 0.628 | 1.610 (1,31) | 0.214 | 2.820 (1,34) | 0.103 | 0.175 (1,34) | 0.679 |
| Right Te1.0 | 0.097 (1,31) | 0.758 | 1.180 (1,31) | 0.286 | 1.200 (1,34) | 0.281 | <.001 (1,34) | 0.994 |
| Left pSTC | 0.401 (1,31) | 0.531 | 2.140 (1,31) | 0.154 | 2.340 (1,34) | 0.135 | 0.718 (1,34) | 0.403 |
| Right pSTC | 0.644 (1,31) | 0.428 | 0.075 (1,31) | 0.787 | 0.805 (1,34) | 0.376 | 0.086 (1,34) | 0.771 |

*Note.* Type 2 Sums of Squares. Factors in the analysis were: condition (HED, LED), hearing test results (PTA or SRT), and age. Only results from the covariates are shown.

